# Are marine biodiversity hotspots still blackspots for barcoding?

**DOI:** 10.1101/2021.07.12.448298

**Authors:** Francesco Mugnai, Emese Meglécz, CoMBoMed group, Federica Costantini, Marco Abbiati, Giorgio Bavestrello, Fabio Bertasi, Marzia Bo, María Capa, Anne Chenuil, Marina Antonia Colangelo, Olivier De Clerck, José Miguel Gutiérrez, Loretta Lattanzi, Michèle Leduc, Daniel Martin, Kenan Oguz Matterson, Barbara Mikac, Laetitia Plaisance, Massimo Ponti, Ana Riesgo, Vincent Rossi, Eva Turicchia, Andrea Waeschenbach, Owen S Wangensteen

## Abstract

Marine biodiversity underpins ecosystem health and societal well-being. Preservation of biodiversity hotspots is a global challenge. Molecular tools, like DNA barcoding and metabarcoding, hold great potential for biodiversity monitoring, possibly outperforming more traditional taxonomic methods. However, metabarcoding-based biodiversity assessments are limited by the availability of sequences in barcoding reference databases; a lack thereof results in high percentages of unassigned sequences. In this study we (i) present the current status of known vs. barcoded marine species at a global scale based on online taxonomic and genetic databases; and (ii) compare the current status with data from ten years ago. Then we analyzed occurrence data of marine animal species from five Large Marine Ecosystems (LMEs) classified as biodiversity hotspots, to identify any consistent disparities in COI barcoding coverage between geographic regions and at phylum level. Barcoding coverage varied among LMEs (from 36.8% to 62.4% COI-barcoded species) and phyla (from 4.8% to 74.7% COI-barcoded species), with Porifera, Bryozoa and Platyhelminthes being highly underrepresented, compared to Chordata, Arthropoda and Mollusca. We demonstrate that although barcoded marine species increased from 9.5% to 14.2% since the last assessment in 2011, about 15,000 (corresponding to 7.8% increase) new species were described from 2011 to 2021. The next ten years will thus be crucial to enroll concrete collaborative measures and long term initiatives (e.g., Horizon 2030, Ocean Decade) to populate barcoding libraries for the marine realm.

---

Healthy and well-functioning ecosystems are crucial for providing essential goods and services (e.g., primary production, climate regulation, CO_2_ sequestration). A critical ecohealth indicator (*sensu* Charron 2012) is biodiversity (Myers et al. 2000) as its loss disrupts ecosystem processes and services. Unfortunately, global biodiversity is predicted to decline over the 21^st^ century due to anthropogenic stressors such as commercial species overexploitation, habitat artificialization and destruction, eutrophication, pollution, and introduction of non-indigenous species (NIS) (Pereira et al. 2010). These pressures intensify the effects of climate change by accentuating associated alterations in oceanic biogeochemistry (Harley 2011) and cause local and global biodiversity losses and spatial reshuffling of species (Pecl et al. 2017; Blowes et al. 2019). Marine biodiversity loss and redistribution are recognized as planetary challenges (Worm et al. 2006; Pimm et al. 2014), but the effective implementation of conservation initiatives depends on having an accurate knowledge of species diversity and their geographic distributions. Therefore, comprehensive biodiversity censuses, particularly of vulnerable marine ecosystems, are imperative to inform policy-making.

Biodiversity studies typically focus on charismatic or commercially-valuable species (Troudet et al. 2017) or well-known biodiversity hotspots (Myers et al. 2000; Jenkins and Van Houtan 2016), where high species richness, endemism and human pressures coincide, leaving many oceanic regions largely unexplored (Snelgrove 2016). While there is a great need to characterize marine biodiversity more widely, few resources are dedicated to taxonomic research and training, which are fundamental for identifying and describing new species (de Carvalho et al. 2007; Boero 2010), particularly of small and cryptic organisms. Furthermore, morphology based biodiversity assessments are time consuming and require highly specialized taxonomic knowledge, which is fast disappearing in the absence of funding. Although genetic characterization (barcoding) of species is highly effective in identifying known taxonomic entities (DeSalle and Goldstein 2019), an entirely molecular approach applied to lesser known taxa or geographic regions goes little beyond identifying molecular operational taxonomic units (MOTUs). Thus, an integrative approach (combining barcoding and morphology) is needed for reliable species identification and high-quality biodiversity assessments.

The rapid generation of high volumes of data resulting from the recent advances in “Omics” technologies and high-throughput sequencing methods poses great opportunities for metabarcoding, metagenomics and metatranscriptomics studies (Raupach et al. 2016; Wangensteen et al. 2018; Holman et al. 2019). Yet, substantial gaps remain in public DNA barcoding databases, leading to a high proportion of unassigned sequences in metabarcoding-based studies (e.g., Martin et al. 2021), and substantial uncertainty in the estimated number of species living in a surveyed area (Valentini et al. 2016). Furthermore, many national and regional initiatives aimed at expanding DNA barcode databases are not providing complete species documentation, including morphology, ecology, and biology. Moreover, databases are usually not fully curated and harmonized by the time funds are exhausted after project completions (Collins et al. 2020). Consequently, some databases may have limited accessibility or are taken offline only a few years after their publication, and the obtained sequences are not always uploaded on public barcoding databases (see Online Resource S1, based on Radulovici et al. 2010 and Trivedi et al. 2016). In addition, most DNA barcoding initiatives were, and still are, focusing on terrestrial or freshwater environments (see Online Resource S1) or on marine organisms that can serve as bioindicators of environmental impacts (Weigand et al., 2019). However, more substantial contributions are needed to improve the genetic knowledge for underrepresented marine taxa at a global scale. Previous attempts of quantifying the proportion of barcoded species across all marine animal species (Bucklin et al. 2011; Bergsten et al. 2012; Vargas et al. 2012; Ratnasingham and Hebert 2013; Aylagas et al. 2018; Weigand et al. 2019; Ramirez et al. 2020) found that only 9.5% of the 192,702 described species had been assigned a molecular cytochrome *c* oxidase subunit I (COI) barcode (Bucklin et al. 2011).

The number of marine species discovered has significantly progressed on a global scale during the last decade (Mora et al. 2011; Costello et al. 2012; Appeltans et al. 2012). Currently, around 238,000 marine species have been described (Horton et al. 2021), with about 2,000 new species added each year (Costello et al. 2012). Here, we provide an updated estimate of COI-barcoded marine animal species at a global scale. Moreover, we selected five Large Marine Ecosystems (LMEs, https://www.lmehub.net) considered as biodiversity hotspots (Mediterranean Sea, Caribbean Sea, North Sea, Indonesian Sea, and Red Sea) to detect possible disparities in COI barcoding coverage at regional scale and to underline where more efforts should be enrolled from now on.

Data for all marine animal species worldwide were gathered from the World Register of Marine Species (WoRMS, Horton et al. 2021), while their presence within the five LMEs was retrieved from the Ocean Biodiversity Information System (OBIS, Intergovernmental Oceanographic Commission of UNESCO; http://www.iobis.org accessed on May 2021). Correspondent availability of COI barcode records at species level were retrieved from the National Center for Biotechnology Information (NCBI, https://www.ncbi.nlm.nih.gov) and Barcode of Life Data System (BOLD, http://www.boldsystems.org) repositories using custom Perl scripts (https://osf.io/qsn5e/).

Globally, from 207,686 known marine animal species, only 14.5% (NCBI) and 13.8% (BOLD) possessed COI barcodes. This represents 4.3% and 5% increases (NCBI and BOLD respectively) (see Online Resource S2) since the estimate by Bucklin et al. (2011). The five LMEs encompass a total of 34,286 nominal marine animal species, with 14,472 (42.2%, NCBI) and 14,351 (41.9%, BOLD) having at least one COI sequence (Fig. 1). Evidently, the proportion of barcoded species in these hotspots largely exceeds that at global level. Nevertheless, the current LMEs barcoding data are still not sufficient for comprehensive biodiversity assessments. Furthermore, this barcoding coverage and species richness varies across LMEs (Fig. 1). The North and Indonesian Seas have comparable species richness, but the number of barcoded species is much higher (+15.8% for both NCBI and BOLD) in the latter. Variable research intensity and disparity in sampling efforts and access to marine biological collections may explain the differences in number and relative occurrences (i.e., frequency) of detected species, as well as the variable barcoding coverage among the LMEs (Collins et al. 2020) (Fig. 2A). In particular, the North Sea is the most intensively studied LME for which species occurrence records started in 1753, while research efforts for the other four LMEs started as late as the 1970s. Moreover, the magnitude of occurrence records does not follow that of species richness. The North Sea has a species richness comparable to the Indonesian Sea (8255 species), but it is considerably lower than that of the Mediterranean (9977 species) and Caribbean (12161 species) Seas. However, it has several millions more species occurrences (9.57 millions) than the other LMEs (Fig. 2B).

**Fig. 1.**
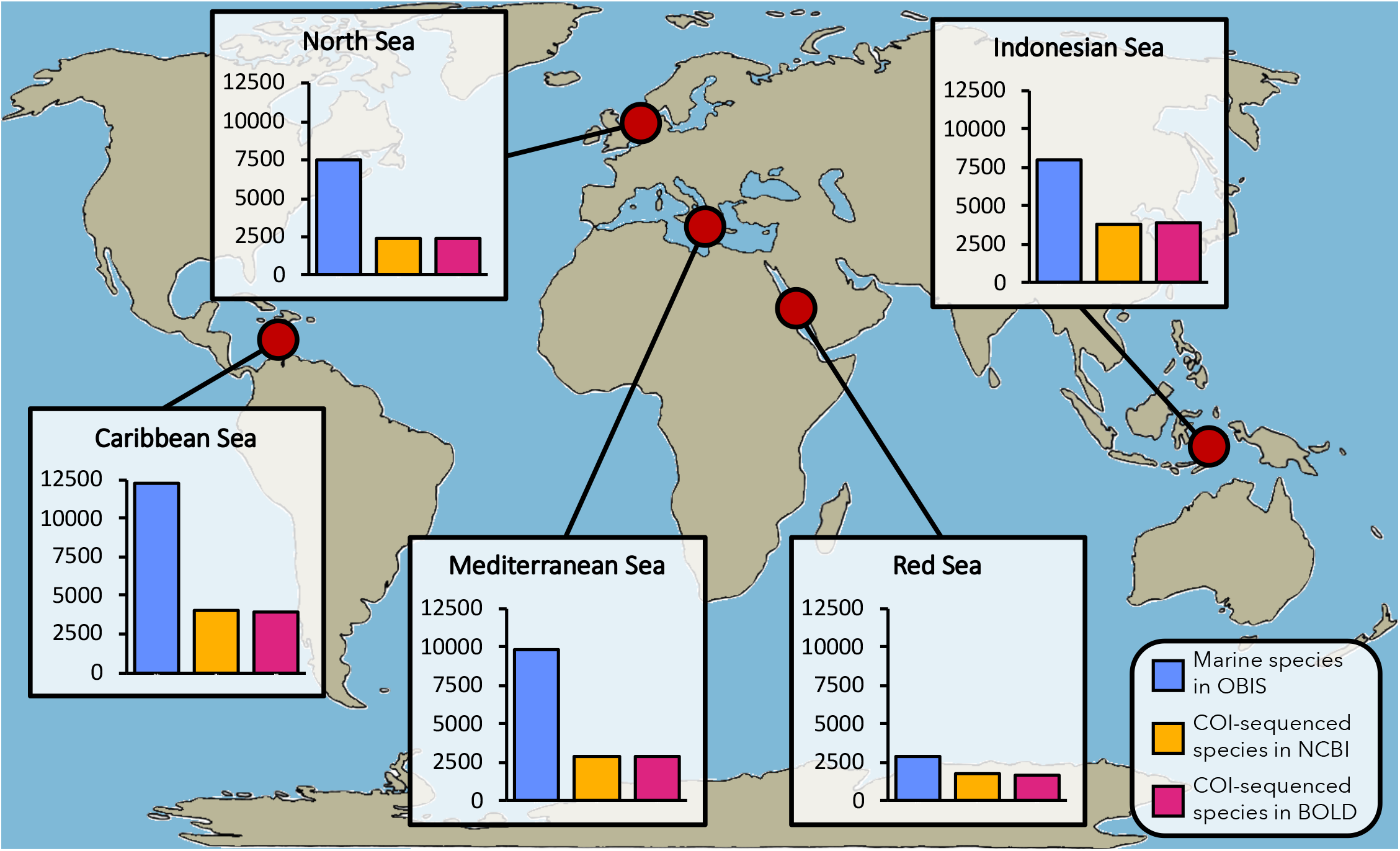
Number of nominal marine and COI-sequenced Animalia species for the five target Large Marine Ecosystems (data accessed in May 2021)

**Fig. 2.**
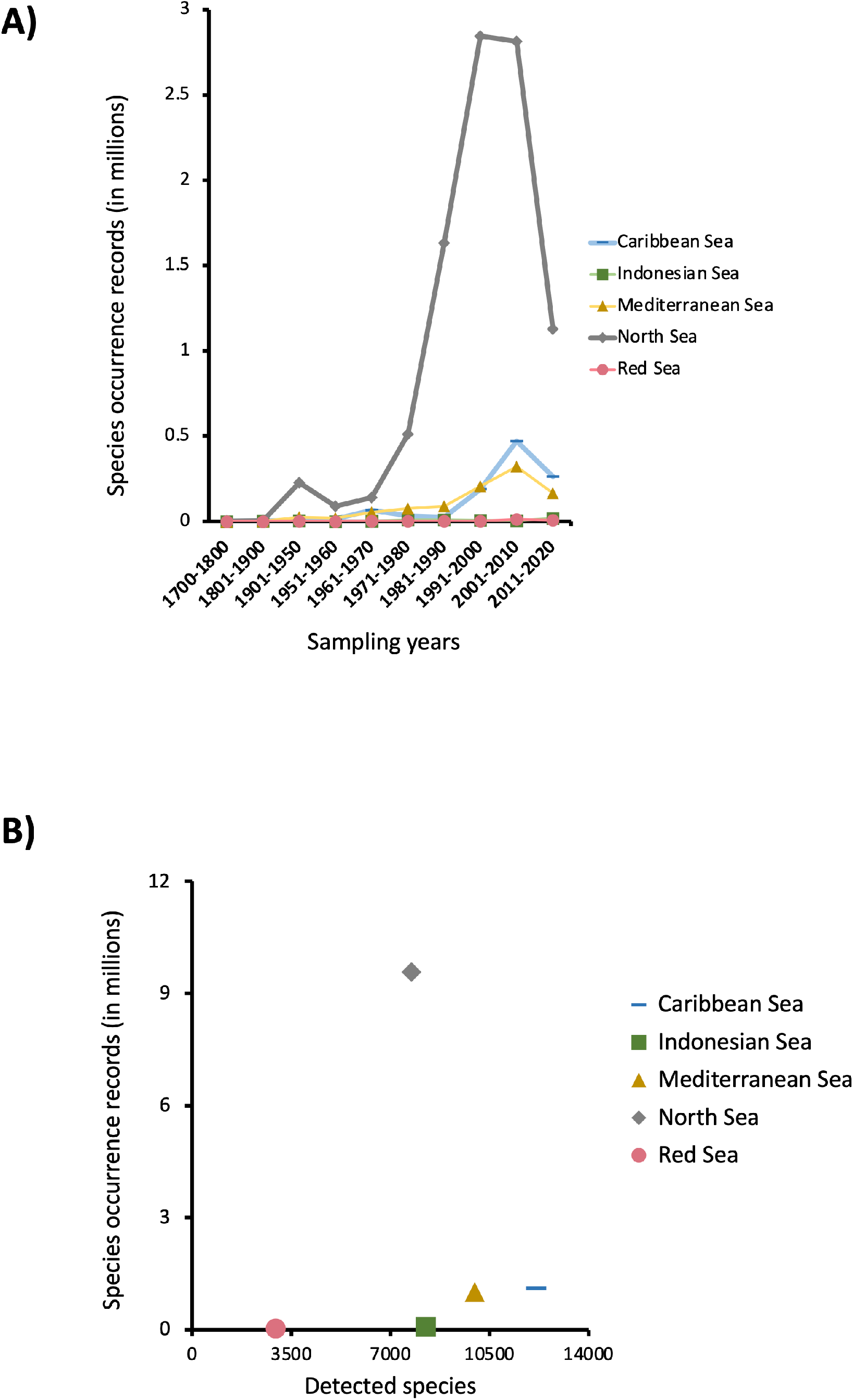
**(a)** Species occurrences over time for each target Large Marine Ecosystem and **(b)** total species occurrences relative to total number of described species per each LME (data accessed in May 2021)

Regarding taxon-specific statistics, the phyla with the highest numbers of nominal and COI-sequenced species in the five LMEs were Chordata, Arthropoda and Mollusca, followed by Annelida and Cnidaria (Fig. 3) (see Online Resource S3 for LME-specific data). COI barcoding coverage for Porifera, Bryozoa, and Platyhelminthes was very low, which is likely due to (1) high crypticity (Trontelj and Fišer 2009; Fehlauer-Ale et al. 2014), (2) variability in the annealing position of traditional COI primers leading to failed PCR amplification, (3) co-amplification of epibionts and endosymbionts (Vargas et al. 2012). Chordata clearly shows the highest number of nominal species and barcoding coverage (68.6% NCBI, 71.5% BOLD). Most of these contributions originate from Vertebrata (70.1% NCBI, 73.1% BOLD), whereas non-vertebrate Chordata (Ascidians) are highly underrepresented (36.3% NCBI, 35.0% BOLD) (see Online Resources S4).

**Fig. 3.**
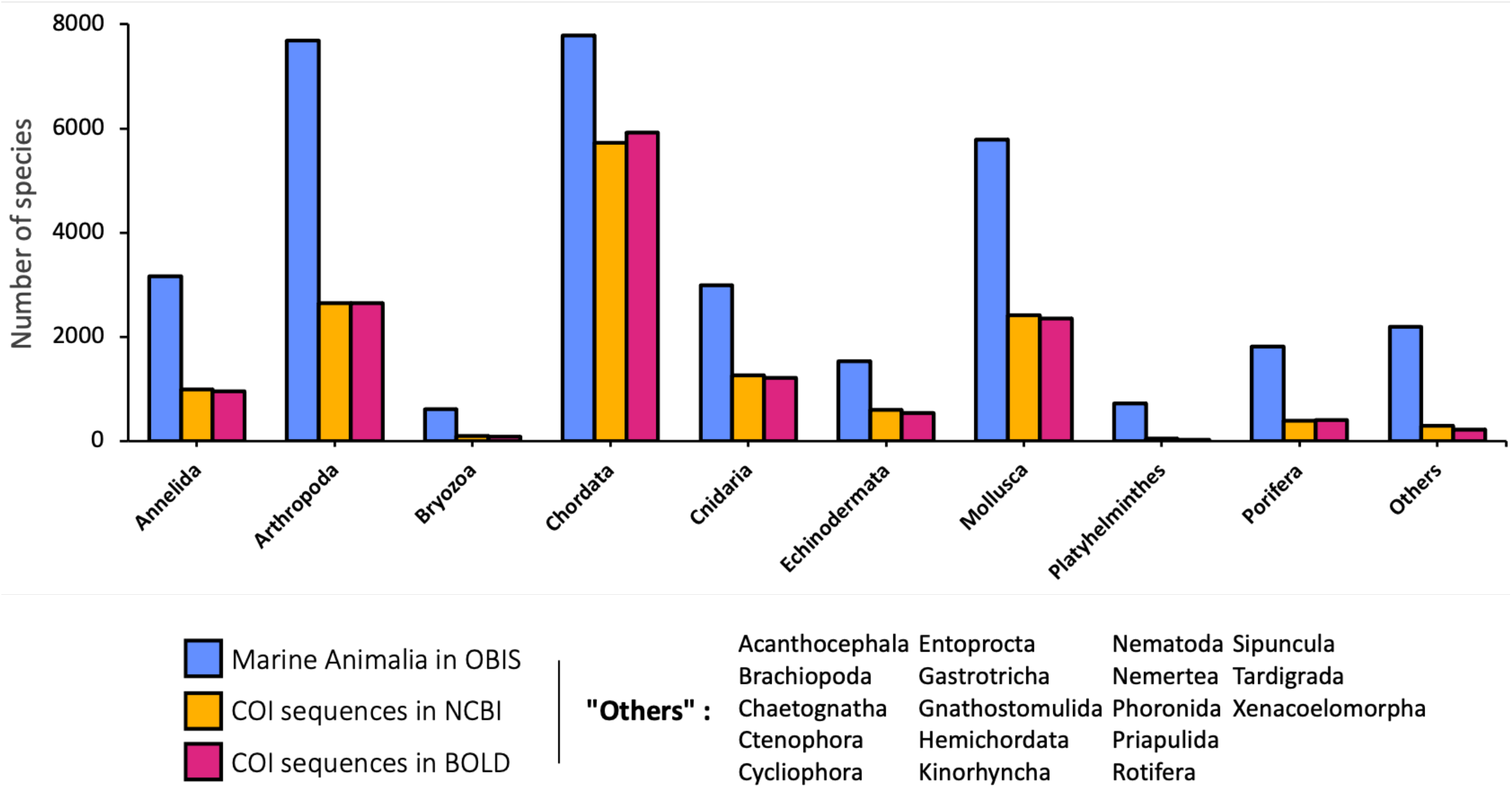
Cumulative number of nominal marine Animalia and COI-sequenced species per phylum in the five target Large Marine Ecosystems (data accessed in May 2021)

Our results evidence a global lack of COI barcode availability for a large proportion of marine invertebrates, with the North, Caribbean, and Mediterranean Seas being highly underrepresented in barcode databases, thus requiring further efforts. The ongoing EU’s key funding programme for research and innovation “Horizon 2030” (2021-2027) allocates 8,953 Bi€ for Pillar II, Cluster 6 “Food, Bioeconomy, Natural Resources, Agriculture & Environment”, offering co-funded initiatives such as “Rescuing Biodiversity to Safeguard Life on Earth” (https://ec.europa.eu/info/horizon-europe_en) that could be pivotal for increasing the biodiversity knowledge at European level, including the marine realm.

To bridge the gaps in biodiversity assessment and monitoring worldwide, national and international collaborations among multidisciplinary teams (including taxonomists, molecular biologists, bioinformaticians, evolutionary biologists, ecologists, and environmental scientists) must be fostered. Such an integrated approach enables physical specimens, accessioned in permanent natural history museum collections, to be linked to molecular, morphological and ecological data, creating meaningful and long-lived reference data repositories. These efforts need to be accompanied by creating an interactive virtual platform able to interface with different database sources, showing up-to-date species-specific ecological, morphological and molecular data. Configuring these public repository following a top-down layout linking other subset databases based on FAIR principles (Findable, Accessible, Interoperable and Reusable, http://wilkinsonlab.info/node/FAIR) is mandatory to allow efficient data sharing.

European initiatives such as DNAquanet (https://dnaqua.net), MBON (https://marinebon.org), SEAMoBB (https://seamobb.osupytheas.fr) are working to reduce morpho-genetic gaps and to expand our knowledge on marine biodiversity. Such initiatives should be further promoted worldwide. The BIOSCAN initiative (2019-2027, https://ibol.org/programs/bioscan/) from the iBOL Consortium aimed at “Revealing species, their interactions, and dynamics” is expected to promote new species descriptions and molecular characterization of terrestrial, freshwater, and marine species. Last but not least, in the present era of unprecedented marine biodiversity loss, society as a whole must be enthused about marine life in view to support measures that preserve marine biodiversity at a global scale. To attain this goal, citizen science initiatives (e.g., marine BioBlitz - https://www.abol.ac.at/en/abol-bioblitz-2020/, LifeWatch - https://www.lifewatch.eu, educational field activities etc.) will be pivotal, as emphasized by the UN Decade of Ocean Science for Sustainable Development (https://www.oceandecade.org).

## Acknowledgements

This paper is a result of the CoMBoMed workshop (Census of Marine Mediterranean Benthic Biodiversity: an integrative metabarcoding approach), held in Ravenna (Italy) on February 11-14, 2019 and organized by the Department of Biological, Geological and Environmental Sciences (BiGeA) of the University of Bologna (UniBo). The CoMBoMed initiative was supported by the European Marine Research Network (EUROMARINE Network), the Inter-Departmental Research Centre for Environmental Sciences (CIRSA – UniBo), the Cultural Heritage Department (DBC - UniBo, https://beniculturali.unibo.it/it), the Fondazione Flaminia and the ERANet Mar-Tera Project SEAMoBB (Solutions for sEmi-Automated Monitoring of Benthic Biodiversity).

