## Supplemental Material for "Are marine biodiversity hotspots still blackspots for barcoding?"

### **Online Resource S1**: DNA barcoding initiatives

| Initiative | Old website | Started in | Ended in | Active until | Still curated | Source | Actual website |
|---|---|---|---|---|---|---|---|
| The Canadian Barcode of Life Network | www.BOLNET.ca | 2005 | 2016 | 2016 | no | | www.nserc-crsng.gc.ca/Business-Entreprise/How-Comment/Networks-Reseaux/DNABarcode-CodeABarreADN_eng.asp |
| European Barcode Of Life (EUBOL) | www.ecbol.org/ | 2008 (?) | 2016 | 2016 | no | | www.bioscaneurope.org |
| Norwegian Barcode Of Life (NorBOL) | www.dnabarcoding.no/en/ | 2007 | 2016 | 2016 | no | | www.norbol.org/en/ |
| Mexican Barcode Of Life (MexBOL) | www.mexbol.org | 2013 (?) | 2016 | 2016 | no | Trivedi et al. 2016 | www.mexbol.com.mx |
| Japanese Barcode Of Life (JBOLI) | www.jboli.org/ | 2007 | | now | unknown | | |
| Human health Barcode Of Life (HealthBOL) | not found | 2010 (?) | | unknown | unknown | | |
| Polar life Barcode Of Life (PolarBOL) | not found | | | unknown | unknown | | |
| Quarantine and plant pathogens (QBOL, as part of the ECBOL) | www.qbol.org/en/qbol.htm | 2009 | | now | unknown | | |
| Formicidae Barcode Of Life | www.formicidaebol.org/ | 2008 | 2015 | unknown | no | | |
| All Birds Barcoding Initiative (ABBI) | www.barcodingbirds.org/ | 2005 | 2015 | unknown | no | | |
| Trichoptera Barcode Of Life | www.trichopterabol.org/ | | 2015 | unknown | no | | |
| Fish Barcode Of Life (FISH-BOL) | www.fishbol.org/ | 2004 | 2015 | unknown | no | | |
| Lepidoptera Barcode Of Life | www.lepbarcoding.org/ | | 2015 | unknown | no | Barcoding 500K Initiative (2010-2015): www.ibol.org/phase1/about-us/campaigns/ | |
| Mammal Barcode Of Life | www.mammaliabol.org/ | | 2015 | unknown | no | | |
| Mosquito Barcoding Initiative | not found | | 2015 | unknown | unknown | | |
| Marine Barcode Of Life | www.marinebarcoding.org/ | 2009 (?) | 2015 | unknown | unknown | | |
| Sponge Barcode project (SpongeBOL) | www.spongebarcoding.org | 2018 (?) | 2015 | now | yes | | |
| Bee Barcode of Life Initiative (Bee-BOL) | not found | 2008 (?) | 2015 | unknown | unknown | | |
| Skatebase | www.skatebase.org | 2014 | | now | no | | |
| Census of Coral Reefs Ecosystems (CReefs) | www.origin-apps-pifsc.fisheries.noaa.gov/cred/creefs.php | 2006 (?) | 2010 | now | unknown | | |
| Census of Marine Zooplankton (CMarZ) | www.cmarz.org | 2004 | 2010 | 2011 | no | | |
| Census of Antarctic Marine Life (CAML) | www.antarctica.gov.au/news/2009/australian-polar-research-at-close-of-ipy/census-of-antarctic-marine-life/ | 2005 | | now (?) | yes (?) | Census of Marine Life: www.coml.org | |
| International Census of Marine Microbes (ICoMM) | www.icomm.mbl.edu | | 2010 | | no | | |

**Online Resource S2**: Morphological and molecular data retrieved in May 2021 and compared with Bucklin et al., 2011

| Our data - 22/05/2021 | | | | | | Our data - 22/05/2021: NCBI – BOLD mean | | | | Bucklin et al., 2011 data | | | | Taxonomy changes |
|---|---|---|---|---|---|---|---|---|---|---|---|---|---|---|
| **Phylum** | **Species** | **NCBI** | **BOLD** | **NCBI%** | **BOLD%** | **Phylum** | **Species** | **Barcoded** | **Barcoded%** | **Phylum** | **Species** | **Barcoded** | **Barcoded%** | |
| Acanthocephala | 517 | 80 | 45 | 15.5 | 8.7 | Acanthocephala | 517 | 63 | 12.2 | Acanthocephala | 600 | 10 | 1.7 | |
| Annelida | 13812 | 1630 | 1466 | 11.8 | 10.6 | Annelida | 13812 | 1548 | 11.2 | Annelida | 12324 | 637 | 5.2 | Echiura |
| Arthropoda | 58492 | 6203 | 5822 | 10.6 | 10 | Arthropoda | 58492 | 6013 | 10.3 | Arthropoda | 47217 | 3580 | 7.6 | |
| Brachiopoda | 416 | 32 | 31 | 7.7 | 7.5 | Brachiopoda | 416 | 32 | 7.7 | Brachiopoda | 550 | 35 | 6.4 | |
| Bryozoa | 6381 | 187 | 164 | 2.9 | 2.6 | Bryozoa | 6381 | 176 | 2.8 | Bryozoa | 5700 | 20 | 0.4 | |
| Chaetognatha | 134 | 23 | 25 | 17.2 | 18.7 | Chaetognatha | 134 | 24 | 17.9 | Chaetognatha | 121 | 23 | 19 | |
| Chordata | 24229 | 10225 | 10790 | 42.2 | 44.5 | Chordata | 24229 | 10508 | 43.4 | Chordata | 21517 | 7279 | 33.8 | |
| Cnidaria | 12098 | 1745 | 1637 | 14.4 | 13.5 | Cnidaria | 12098 | 1691 | 14 | Cnidaria | 9795 | 594 | 6.1 | |
| Ctenophora | 210 | 16 | 12 | 7.6 | 5.7 | Ctenophora | 210 | 14 | 6.7 | Ctenophora | 166 | 0 | 0 | |
| Cycliophora | 2 | 2 | 2 | 100 | 100 | Cycliophora | 2 | 2 | 100 | Cycliophora | 1 | 1 | 100 | |
| Dicyemida | 122 | 11 | 0 | 9 | 0 | Dicyemida | 122 | 6 | 4.9 | Dicyemida | 82 | 0 | 0 | Rhombozoa |
| Echinodermata | 8019 | 1512 | 1329 | 18.9 | 16.6 | Echinodermata | 8019 | 1421 | 17.7 | Echinodermata | 7000 | 771 | 11 | |
| Entoprocta | 188 | 14 | 4 | 7.4 | 2.1 | Entoprocta | 188 | 9 | 4.8 | Entoprocta | 170 | 0 | 0 | |
| Gastrotricha | 521 | 45 | 42 | 8.6 | 8.1 | Gastrotricha | 521 | 44 | 8.4 | Gastrotricha | 400 | 0 | 0 | |
| Gnathostomulida | 100 | 8 | 7 | 8 | 7 | Gnathostomulida | 100 | 8 | 8 | Gnathostomulida | 97 | 8 | 8.2 | |
| Hemichordata | 132 | 2 | 9 | 1.5 | 6.8 | Hemichordata | 132 | 6 | 4.5 | Hemichordata | 106 | 2 | 1.9 | |
| Kinorhyncha | 307 | 27 | 24 | 8.8 | 7.8 | Kinorhyncha | 307 | 26 | 8.5 | Kinorhyncha | 130 | 0 | 0 | |
| Loricifera | 29 | 0 | 0 | 0 | 0 | Loricifera | 29 | 0 | 0 | Loricifera | 18 | 0 | 0 | |
| Mollusca | 50841 | 6568 | 5847 | 12.9 | 11.5 | Mollusca | 50841 | 6208 | 12.2 | Mollusca | 52525 | 4813 | 9.2 | |
| Nematoda | 6465 | 177 | 122 | 2.7 | 1.9 | Nematoda | 6465 | 150 | 2.3 | Nematoda | 12000 | 180 | 1.5 | |
| Nematomorpha | 5 | 0 | 0 | 0 | 0 | Nematomorpha | 5 | 0 | 0 | Nematomorpha | 5 | 0 | 0 | |
| Nemertea | 1312 | 220 | 204 | 16.8 | 15.5 | Nemertea | 1312 | 212 | 16.2 | Nemertea | 1230 | 81 | 6.6 | |
| Orthonectida | 26 | 0 | 0 | 0 | 0 | Orthonectida | 26 | 0 | 0 | Orthonectida | 24 | 0 | 0 | |
| Phoronida | 13 | 10 | 9 | 76.9 | 69.2 | Phoronida | 13 | 10 | 76.9 | Phoronida | 10 | 0 | 0 | |
| Placozoa | 3 | 0 | 2 | 0 | 66.7 | Placozoa | 3 | 1 | 33.3 | Here were Rhombozoa (now Dicyemida) | | | | |
| Platyhelminthes | 12894 | 520 | 181 | 4 | 1.4 | Platyhelminthes | 12894 | 351 | 2.7 | Platyhelminthes | 15000 | 124 | 0.8 | |
| Porifera | 9314 | 736 | 721 | 7.9 | 7.7 | Porifera | 9314 | 729 | 7.8 | Porifera | 5500 | 67 | 1.2 | |
| Priapulida | 22 | 4 | 3 | 18.2 | 13.6 | Priapulida | 22 | 4 | 18.2 | Priapulida | 8 | 1 | 12.5 | |
| Rotifera | 192 | 42 | 28 | 21.9 | 14.6 | Rotifera | 192 | 35 | 18.2 | Rotifera | 50 | 20 | 40 | |
| Sipuncula | 201 | 51 | 50 | 25.4 | 24.9 | Sipuncula | 201 | 51 | 25.4 | Sipuncula | 144 | 15 | 10.4 | |
| Tardigrada | 230 | 5 | 4 | 2.2 | 1.7 | Tardigrada | 230 | 5 | 2.2 | Tardigrada | 212 | 9 | 4.2 | |
| Xenacoelomorpha | 459 | 66 | 62 | 14.4 | 13.5 | Xenacoelomorpha | 459 | 64 | 13.9 | | | | | |
| TOTALS | 207686 | 30161 | 28642 | 14.5 | 13.8 | TOTALS | 207686 | 29411 | 14.2 | TOTALS | 192702 | 18270 | 9.5 | |

**Online Resource S3**: Number of nominal marine Animalia species and relative COI-sequenced taxa per phylum in the five target Large Marine Ecosystems

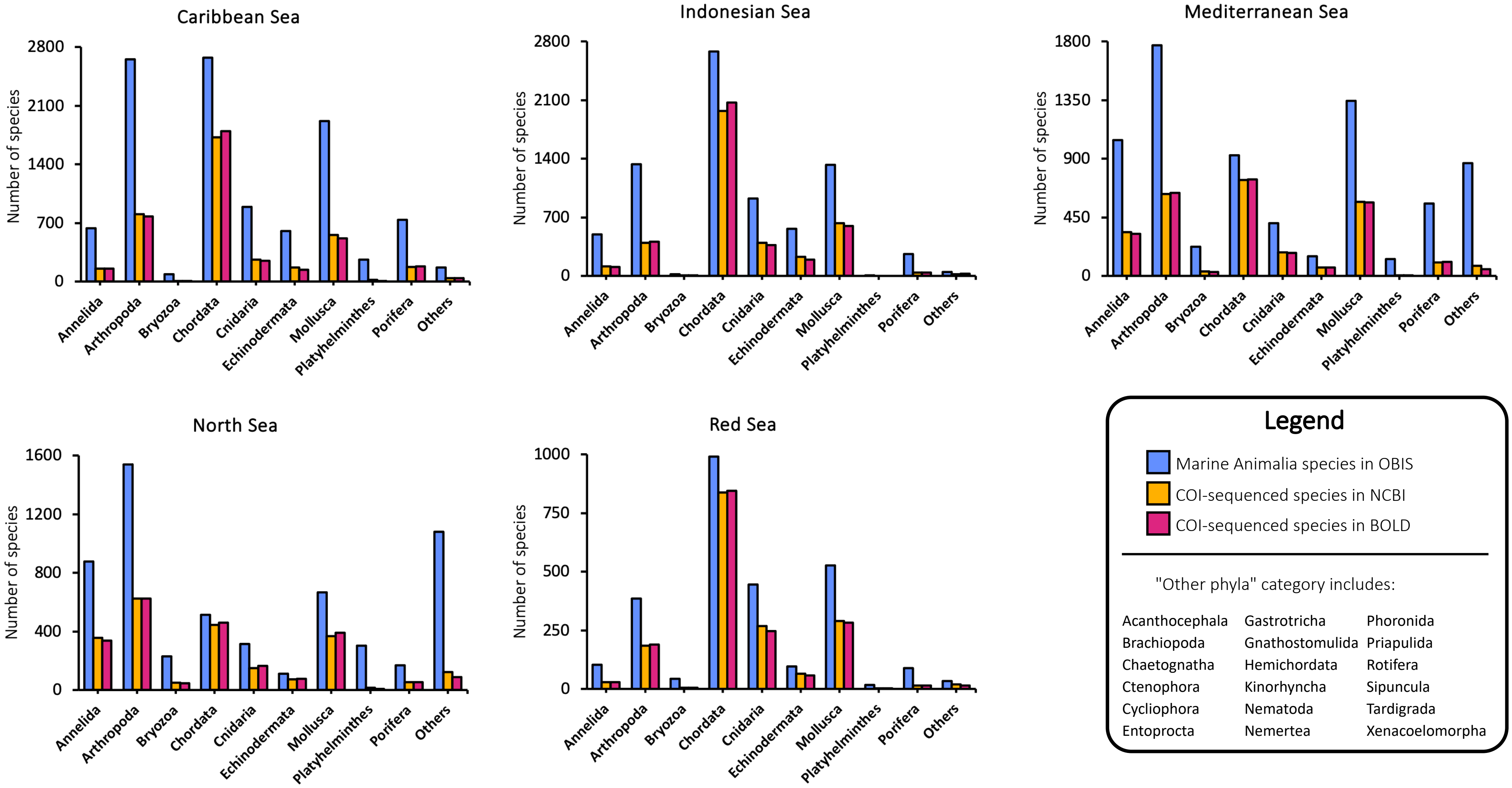

**Online Resource S4**: **A)** Total number of nominal marine Chordata species and relative COI-sequenced taxa per phylum and **B)** relative values scattered for each of the five target Large Marine Ecosystems

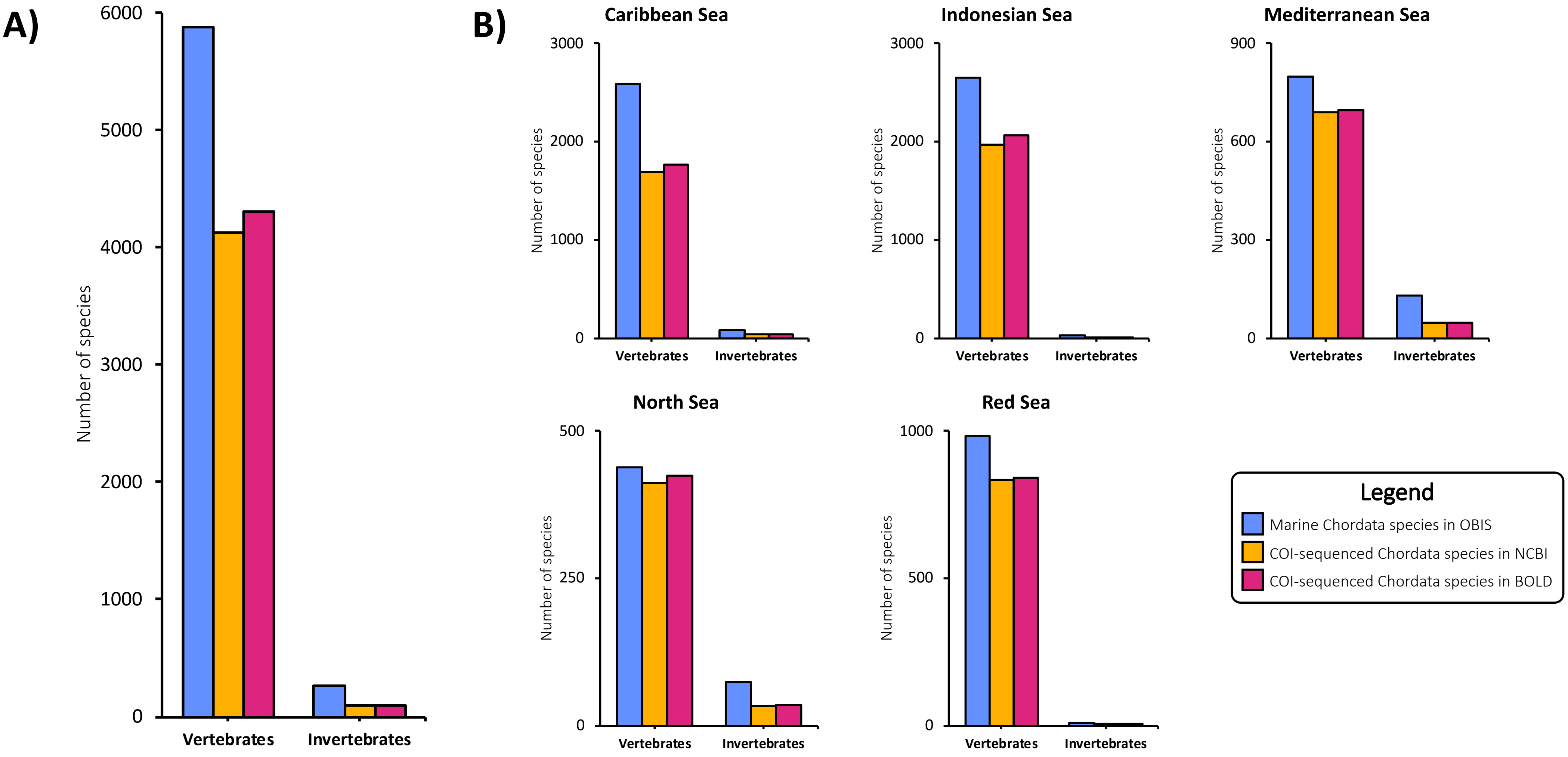
